# Random Forest Feature Selection, Fusion and Ensemble Strategy: Combining Multiple Morphological MRI Measures to Discriminate among healthy elderly, MCI, cMCI and Alzheimer's disease patients: from the Alzheimer’s disease neuroimaging initiative (ADNI) database

**DOI:** 10.1101/236141

**Authors:** S.I. Dimitriadis, D. Liparas, Magda N. Tsolaki

**Affiliations:** Neuroscience and Mental Health Research Institute, Cardiff University, Cardiff, UK.; Cardiff University Brain Research Imaging Centre (CUBRIC), School of Psychology, Cardiff University, Cardiff, UK; MRC Centre for Neuropsychiatric Genetics and Genomics, Institute of Psychological Medicine and Clinical Neurosciences, Cardiff School of Medicine, Cardiff University, Cardiff, UK; Neuroinformatics Group, (CUBRIC), School of Psychology, Cardiff University, Cardiff, UK; School of Psychology, Cardiff University, Cardiff, UK; High Performance Computing Center Stuttgart (HLRS), University of Stuttgart, Stuttgart, Germany; Department of Informatics, Aristotle University of Thessaloniki, Thessaloniki, Greece; 3rd Department of Neurology, Medical School, Aristotle University of Thessaloniki, Thessaloniki, Greece

**Author notes:** The first two authors contributed equally. All the data used in preparation of this article were obtained from the Alzheimer's Disease Neuroimaging Initiative (ADNI) database (adni.loni.usc.edu). The investigators within the ADNI contributed to the design and implementation of ADNI and/or provided data but did not participate in analysis or writing of this report. A complete listing of ADNI investigators can be found at: http://adni.loni.usc.edu/wp-content/uploads/how_to_apply/ADNI_Acknowledgement_List.pdf. The preprocessing of the T1-weighted Magnetic Resonance Images (MRI) was conducted by the organizers of the competition; information can be found here: https://inclass.kaggle.com/c/mci-prediction.

## Abstract

**Background:** In the era of computer-assisted diagnostic tools for various brain diseases, Alzheimer’s disease (AD) covers a large percentage of neuroimaging research, with the main scope being its use in daily practice. However, there has been no study attempting to simultaneously discriminate among Healthy Controls (HC), early mild cognitive impairment (MCI), late MCI (cMCI) and stable AD, using features derived from a single modality, namely MRI.

**New Method:** Based on preprocessed MRI images from the organizers of a neuroimaging challenge^2^, we attempted to quantify the prediction accuracy of multiple morphological MRI features to simultaneously discriminate among HC, MCI, cMCI and AD. We explored the efficacy of a novel scheme that includes multiple feature selections via Random Forest from subsets of the whole set of features (e.g. whole set, left/right hemisphere etc.), Random Forest classification using a fusion approach and ensemble classification via majority voting.

From the ADNI database, 60 HC, 60 MCI, 60 cMCI and 60 AD were used as a training set with known labels. An extra dataset of 160 subjects (HC: 40, MCI: 40, cMCI: 40 and AD: 40) was used as an external blind validation dataset to evaluate the proposed machine learning scheme.

**Results:** In the second blind dataset, we succeeded in a four-class classification of 61.9% by combining MRI-based features with a Random Forest-based Ensemble Strategy. We achieved the best classification accuracy of all teams that participated in this neuroimaging competition.

**Comparison with Existing Method(s):** The results demonstrate the effectiveness of the proposed scheme to simultaneously discriminate among four groups using morphological MRI features for the very first time in the literature.

**Conclusions:** Hence, the proposed machine learning scheme can be used to define single and multi-modal biomarkers for AD.

**HIGHLIGHTS:**

- 1^st^ place in International Challenge for Automated Prediction of MCI from MRI Data
- Multi-class classification of normal control, MCI, converting MCI, and Alzheimer’s disease
- Morphometric measures from 3D T1 brain MRI images have been analysed (ADNI1 cohort).
- A **Random Forest Feature Selection, Fusion and Ensemble Strategy** was applied to classification and prediction of AD.
- Accuracy and robustness have been assessed in a blind dataset

## 1 Introduction

In the era of computer-assisted diagnostic tools for various brain diseases, Alzheimer’s disease (AD) covers a large percentage of neuroimaging research that is applied in daily practice. Pattern recognition approaches, tailored to neuroimaging, offer to the neuroscience community a potential diagnostic tool, particularly though not restricted to magnetic resonance imaging (MRI), which has shown its effectiveness in the diagnosis of AD (O’Brien, 2007).

In recent years and with the release of free available databases, a large number of studies have introduced the use of pattern recognition and machine learning approaches, based on MRI, for the early detection of AD (Gray et al., 2013; Lebedev et al., 2014; Schwarz et al., 2016; Vos et al., 2016; Kalin et al., 2017). The obvious advantage of the proposed semi- or fully-automated methods, compared to visual inspection of an MRI by an expert, e.g. a radiologist or neurologist, is its avoidance of biases and, in many cases, errors to which human diagnosis is subject. All these techniques further improved classification accuracy, primarily through enhancing the stability of the algorithmic pipelines and finally through development of a standardized computerized decision support system, a rapidly progressing field in the overlapping areas of radiology and machine learning (Stivaros et al., 2010; Belle et al., 2013).

Alzheimer’s disease (AD) is a neurogenerative disorder with chronic symptoms that mostly starts slowly and progresses faster over an individual’s lifetime (Mendez, 2012). AD is the main cause of 60% to 70% of dementia, and the first symptom is difficulty in remembering recent events, which is called short-term memory loss (Jahn, 2013). The basic anatomical alteration of AD is hippocampal atrophy, which is highly used as a clinical biomarker (Morra et al., 2009). Complementary to hippocampal atrophy, the atrophy of grey matter in AD could be extended also to the medial temporal lobe and other subcortical structures (Seeley et al., 2009). The localization and the extension of grey matter atrophy can be realized via the visualization of anatomical magnetic resonance imaging (MRI) scans, which is the most commonly used approach for the clinical diagnosis of AD (Frisoni et al., 2010). Previous studies have revealed that AD patients have different volumes and shapes of hippocampi areas, compared to elderly individuals with normal cognitive abilities (Scher et al., 2007). Another study demonstrated that the volumes of the thalamus and the putamen are also reduced in AD (de Jong et al., 2008). Additionally, extended grey matter atrophy has been revealed in AD patients compared to age-matched controls (Karas et al., 2003) and also reduced cortical thickness in another study (Lerch et al., 2005).

In contrast, many significant observations based on MRIs of AD patients with direct comparison with controls cannot be used in a straightforward way in a daily clinical setting. Some of the candidate AD biomarkers have also been detected in healthy aging. Potential MRI-based AD biomarkers are only useful if they can discriminate AD subjects from non-affected subjects at the subject level (Salat et al., 1999). Lately, research on anatomical MRI markers tailored to AD has shifted from group differences to disease detection (de Vos et al., 2016). Measurements of the shape and volume of hippocampi sub-areas and cortical thickness, and voxel-based morphometry (VBM), have been extensively used to separate controls from AD patients with a high range of classification accuracy (Cuingnet et al., 2011; Davatzikos et al.,2011; Querbes et al., 2009; de Vos et al., 2016; Kalin et al., 2017).

Alternative anatomical MRI-based estimates have been mainly used to discriminate AD patients from healthy controls. The principle behind the strategy of using features from different feature sets is that they encapsulate complementary information and their combination could further increase the classification accuracy of AD. VBM and voxel-based cortical thickness demonstrate complementary aspects of age-based decline of grey matter (Hutton et al., 2009). Combining alternative anatomical MRI-based estimates further improved AD classification accuracy (Bron et al., 2015; Wolz et al., 2011; Westman et al., 2013; de Vos et al., 2016; Kalin et al., 2017). Apart from separating AD patients from individuals undergoing healthy aging, these features have proved useful for separating mild cognitive impairment (MCI) converters from individuals with stable MCI (Dyrba et al., 2015a,b; Schouten et al.,2016; Trzepacz et al., 2014; Teipel et al., 2010). Based on the aforementioned studies, it would be very important to demonstrate the effectiveness of combining different but complementary sets of features derived from anatomical MRI to simultaneously discriminate among healthy controls, AD, MCI and cMCI.

We analyzed the T1-weighted Magnetic Resonance Images (MRI) preprocessed by the organizers of a Neuroimaging Challenge/Competition, information on which can be found here: https://inclass.kaggle.com/c/mci-prediction. The anatomical MRI scans were derived from four groups (healthy controls, AD, MCI and cMCI). The MRI features that were extracted are the following: (i) cortical thickness, (ii) cortical surface area, (iii) cortical curvature, (iv) grey matter density, (v) the volume of the cortical and subcortical structures, (vi) the shape of the hippocampus and (v) hippocampal subfield volumes. We attempted to achieve high generalization of the proposed feature extraction and classification strategy with the use of a training set aiming at increasing classification accuracy in a second blind dataset.

In this study, Random Forest (RF) was adopted as a proper ensemble learning algorithm, which is a combination of tree predictors where each tree is built on a random vector built in with features sampled independently and with the same distribution across trees in the forest (Breiman, 2001). In addition, out-of-bag error estimation (OOB) was adopted in order to accurately detect the generalization error of the model. Finally, proximity ratio estimation for late fusion strategies and weighted fusion (Liparas et al., 2014) was applied to further improve the classification accuracy in a blind test set.

We hypothesized that using RF with the complementary techniques that guarantee the generalization of the model based on a training dataset and the incorporation of MRI-based features, it would be possible to achieve a high classification in the aforementioned four-class problem. Additionally, we hypothesized that it is possible to keep the classification performance on the same level as in previous studies, in which three classes were used (HC, MCI and AD).

## 2 Material and Methods

### 2.1 Participants

In particular, MRIs were selected from the **Alzheimer’s disease Neuroimaging Initiative** (ADNI). ADNI is an international project that collects and validates neurological data, such as MRI and PET images, genetics or cognitive tests.

We randomly and automatically selected subjects with a static seed, by using the data analytics platform Konstanz Information Miner (KNIME).

Subjects from ADNI were selected by filtering text files downloaded from the website in three steps. In particular, we used the file containing the conversion of diagnosis for first choosing healthy controls (HC), Alzheimer's patients (AD) and Mild Cognitive Impairment (MCI) who did not convert their diagnosis in the follow up. Then, with the same approach, we selected those with MCI who converted to Alzheimer’s (cMCI). The second step was to obtain demographic and clinical parameters at that timepoint, i.e. age, gender and Mini-Mental State Examination score (MMSE). This dataset was grouped by diagnostic criteria, in order to obtain a balanced number of subjects (100) for each of the four classes (HC, AD, MCI, cMCI).

The last step was to obtain the subjects’ MRI scan ID at the baseline from the file MPRAGEMETA.csv (green area in Fig. 1 in Sarica et al., 2014). In particular, we selected the first MPRAGE sequence (no repetition), acquired at 3 Tesla.

**Figure 1:**
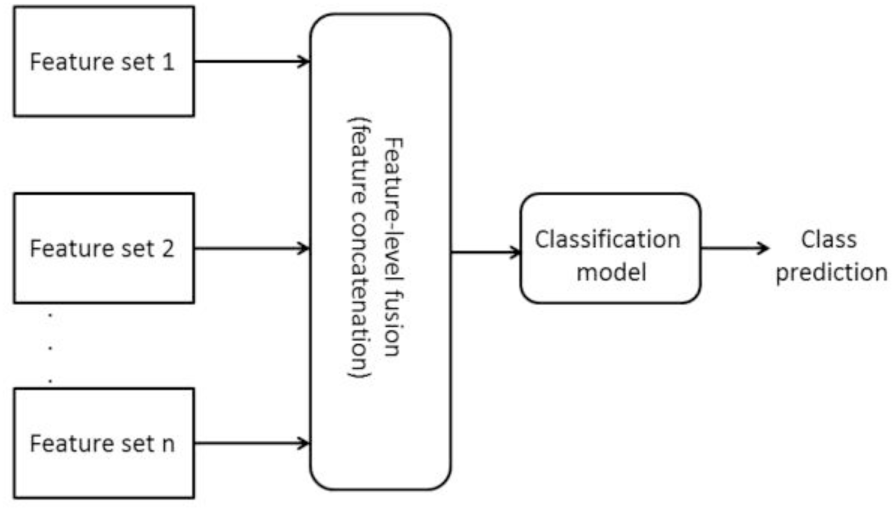
A flowchart that describes the notion of early fusion

Finally, the whole dataset of 400 subjects was split into a training dataset of 240 subjects (60 subjects for each of the four groups) and a testing dataset of 160 subjects (40 subjects for each of the four groups).

Table 1 summarizes the demographics of the training and testing datasets, including the average age, the gender contribution and the average MMSE.

**Table 1.**
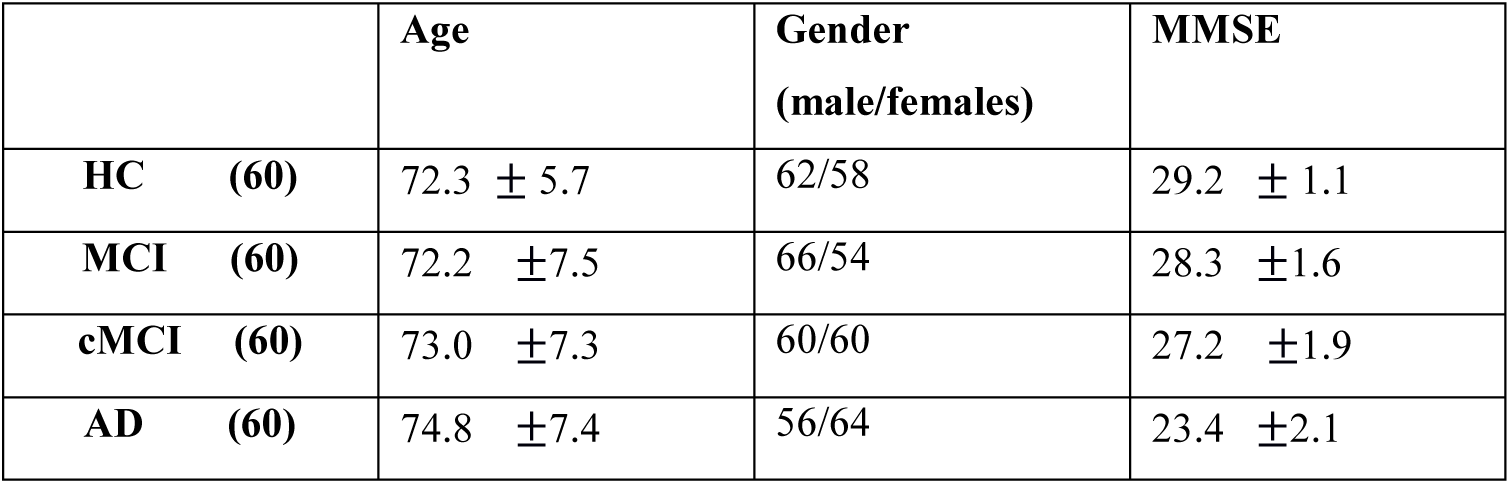
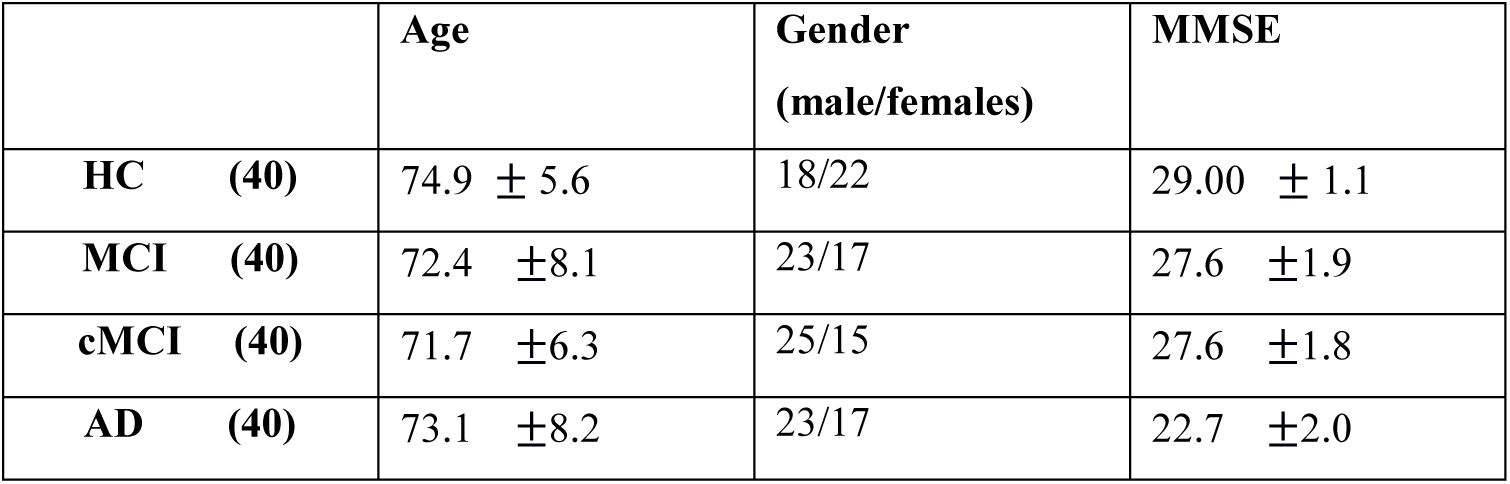
Summary of demographics for the testing dataset

|  | Age | Gender<br>(male/females) | MMSE |
| --- | --- | --- | --- |
| **HC (60)** | 72.3 ± 5.7 | 62/58 | 29.2 ± 1.1 |
| **MCI (60)** | 72.2 ± 7.5 | 66/54 | 28.3 ± 1.6 |
| **cMCI (60)** | 73.0 ± 7.3 | 60/60 | 27.2 ± 1.9 |
| **AD (60)** | 74.8 ± 7.4 | 56/64 | 23.4 ± 2.1 |
| **HC (40)** | 74.9 ± 5.6 | 18/22 | 29.00 ± 1.1 |
| **MCI (40)** | 72.4 ± 8.1 | 23/17 | 27.6 ± 1.9 |
| **cMCI (40)** | 71.7 ± 6.3 | 25/15 | 27.6 ± 1.8 |
| **AD (40)** | 73.1 ± 8.2 | 23/17 | 22.7 ± 2.0 |

### 2.2 MR Image Acquisition

All participants were scanned on a Philips 3 T Achieva MRI scanner. The MRI data acquisition protocol is described in ADNI’s official webpage^2^.

### 2.3 Freesurfer processing and Features Extraction

T1-weighted Magnetic Resonance Images (MRI) were processed by the organizers of the Neuroimaging Challenge/Competition for an automated classification of MCI. Additional information can be found here: https://inclass.kaggle.com/c/mci-prediction.

MRIs were preprocessed by <u>Freesurfer</u> (v5.3), with the standard pipeline (*recon-all-hippo-subfields*) on a computer running GNU/Linux Ubuntu 14.04 with 16 CPUs and 16Gb RAM. We used the KNIME plugin <u>K-Surfer</u> (Sarica et al., 2014) for extracting numerical data produced by Freesurfer into a table format. The organizers of the competition then joined this table with demographical and clinical parameters.

The set of features used for the training procedure are the following:

- MMSE_bl - Mini-mental state examination total score at the baseline of the subject
- Age and
- (i) cortical thickness,(ii) cortical surface area, (iii) cortical curvature, (iv) grey matter density, (v) the volume of the cortical and subcortical structures, (vi) the shape of the hippocampus and (v) Hippocampal subfields volume.

### 2.4 Problem Formulation

The organizers of the International Challenge for Automated Prediction of MCI from MRI Data generated an additional 340 artificial test observations that were joined with the real test observations (4×40=160) in the Challenge test set to form a combined test set of 500 observations. This testing sample was used in the online Kaggle competition platform for the evaluation of the classification performance (Sarica et al., 2016). This set, which can be called an artificial – Challenge dataset, was split into a public and private test set. The competition started online between 21 December 2016 and 1 June 2017 and every team that participated in this neuroimaging competition had the option of one submission per day. After every submission, the organizers returned, via the kaggle web system, the accuracy over 500 subjects, where only 160 subjects were the real blind dataset, while the rest (340 subjects - dummy) were created via a model based on the features from the training dataset. By the end of the challenge on 1 June 2017, the best performance of each team was evaluated and selected based on the private test set. The final evaluation and the ranking of the teams in terms of the classification accuracy was realized based on the Challenge test set which contains the real test data. Finally, the labels of the test data and the confusion matrices were released to the participants and teams that were invited to contribute to this special issue, dedicated to the international challenge for the automated prediction of MCI using MRI data. Our team won the first position in this neuroimaging challenge.

Our best submission was built around an ensemble of five classification models. The construction of these models was based on the well-known Random Forests (RF) machine learning method and its operational capabilities. More specifically, in all models, we performed feature selection using the Gini impurity index, a type of feature importance measurement commonly used in RF. In addition, we employed early fusion, as well as weighted fusion by means of late fusion schemes based on internal mechanisms provided by RF, namely the out-of-bag error and proximity ratios.

In what follows, the theoretical background of the involved methodologies, as well as a description of each classification model that was utilized in our experiments, are provided.

#### 2.4.1 Random Forests

Random Forests (RF) is a popular machine learning method used in classification, regression and other tasks (Breiman, 2001). The methodology involves the construction of a multitude of decision trees and within RF, randomness is employed in the following ways: Firstly, each decision tree is constructed using a different bootstrap sample (a training set that is drawn randomly from the original training data by sampling uniformly and with replacement). Secondly, during the construction of each decision tree, each node split involves the random selection of a subset of *k* variables (from the original variable set), based on which the best split is determined and used. For the prediction of unknown cases, the decisions of the constructed trees are aggregated by employing majority voting for classification and averaging for regression tasks.

The *out-of-bag* (OOB) error estimate is an internal mechanism provided by RF for estimating the generalization error of a constructed model. The OOB error is estimated as follows: RF uses only around 66% (2/3) of the original data in order to build each decision tree.

The other 33% (approximately) of the original data cases, called OOB data, are predicted by the constructed decision tree and are consequently utilized as “test” data. The averaged prediction error for each training case x, using only the trees that do not include *x* in their bootstrap sample, is the OOB error estimate. Additionally, the *proximity matrix* is another useful tool in RF. The way to compute the proximity matrix is the following: for each constructed decision tree in an RF model, all data cases (both training and OOB) are put down that tree. If a pair of cases is found in the same terminal node of the tree, their proximity is increased by one. In this way, a matrix of proximities between all data cases is constructed for the entire RF model. Finally, the proximities in the matrix are normalized by dividing their values by the number of trees in the forest.

Another operational feature of RF is its natural ability to provide a ranking of the importance of variables in a regression or classification problem. This can be achieved in two ways. The first one is based on statistical permutation tests, while the second way, which is used in this study, is based on the *Gini impurity index*. Gini impurity is computed at every node split during the construction of a decision tree in an RF model and is used for measuring the quality of the split in terms of separating the samples of the different classes in the considered node. For a variable, the Gini impurity index is computed as in the following equation:

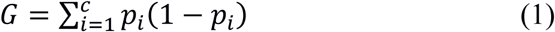

where *c* the number of classes in the variable and *p_i_* the fraction of samples labeled with class *i* in the node.

For a given node split, the values of the Gini impurity index for the two resulting nodes are less than the value for the parent node. If we sum the Gini impurity decreases for each variable in a dataset over all trees in a RF model, we get the corresponding Gini importance measure for each variable, which can consequently be used for feature selection. For more details on the Gini variable importance approach in RF, we refer to (Menze et al., 2009).

#### 2.4.2 Fusion schemes

An interesting and at the same time important challenge in classification tasks is the use of methods for the combination of multiple feature sets (or modalities), a procedure that is known as multimodal fusion. In this context, two basic strategies regarding the level at which fusion is performed can be considered. In the first strategy, known as early fusion, feature-level fusion is performed, where features from the individual feature sets/modalities are concatenated in order to create a common feature vector. Then, a classifier is trained using this common feature vector in order to form the final prediction model. In the second strategy, called late fusion, decision-level fusion is performed, in which a classification model is trained separately for each feature set/modality and the individual results (classifier scores) are fused into a final common decision. The standard way to combine multiple classifiers in late fusion is to compute a weighted sum of the individual classifiers’ scores. Figures 1 and 2 depict the notion of early and late fusion, respectively.

**Figure 2:**
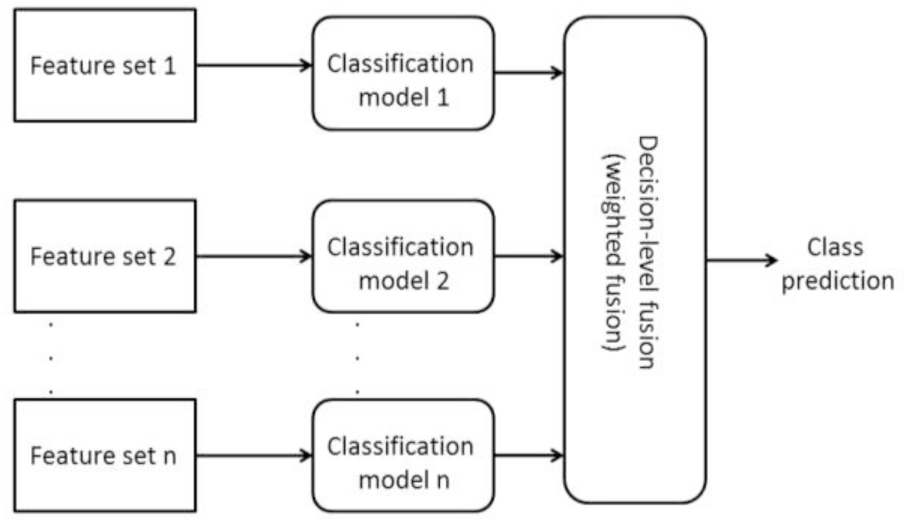
A flowchart that describes the notion of late fusion

In this study, we applied early fusion as well as late fusion strategies based on RF’s operational features, namely the OOB error and proximity ratios (derived from the proximity matrix). The description of these two late fusion strategies is provided below:

Suppose there are two feature sets/modalities, namely *D* and *E*. First, the feature vector from each set is used for training a separate RF model. From the two RF models, the weights for each feature set/modality needs to be computed in order to apply weighted fusion and provide the final RF predictions. The OOB and proximity ratio late fusion strategies are applied as follows:

##### OOB strategy

From the OOB error estimate of each feature set’s RF model, the OOB accuracy values are computed separately for each considered class. These values are then normalized (by dividing them by their sum) and serve as weights for the two feature sets/modalities.

##### Proximity ratio strategy

The same approach, as in the case of the OOB strategy, is followed for the proximity ratio strategy. Nevertheless, instead of utilizing the OOB accuracy values from each RF model, the ratio values between the inner-class and the intra-class proximities (for each class) are used (Zhou et al., 2010). For each RF model, the proximity matrix between all pairs of data cases PR = {*pr_ij_*, i,j = 1,…,n} (n=number of data cases) is constructed, and then the ratio values between the inner-class and the intra-class proximities are computed as in the following equation:

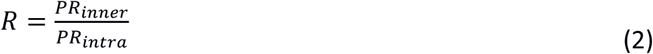

where

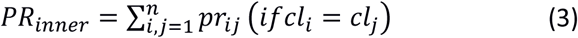

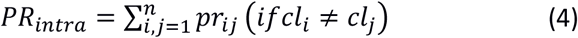

and *cl_i_*, *cl_j_* the class labels of cases *i* and *j*, respectively.

##### Weighted fusion

For the prediction of an unknown case, the RF models provide probability estimates per class for that case. In our example, the probability outputs *P_D_* and *P_E_* from the two feature sets/modalities *D* and *E*, respectively, are multiplied by their corresponding modality weights *W_D_* and *W_E_* (computed either with the OOB strategy or the proximity ratio strategy) and summed in order to produce the final RF predictions as in the following equation:

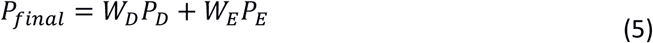

For more details on the aforementioned late fusion strategies, we refer to (Liparas et al., 2014).

#### 2.4.3 Models description

In this section, the features of the five classification models of our best submission’s ensemble are described.

1. Model 1: The first model involved the training of a RF classifier on the whole feature set, as well as feature selection by means of the Gini importance measure, which provided the final feature subset that was used for retraining the RF model.
2. Model 2: In the second model, the following steps were involved:

A. The initially provided feature space was first split into two modalities – feature sets, with each set containing features/measurements from the left or right hemisphere, respectively.
B. A RF model was trained for each modality and, as in the case of Model 1, the Gini importance measure was utilized for selecting the most important features from each modality.
C. The RF models were retrained with the use of the resulting feature subsets.
D. For formulating the final predictions/probability scores from the two RF models, weighted fusion was applied with the use of the proximity ratio late fusion strategy (described in Section 2.4.2).
3. Model 3: With respect to the third model, the exact same approach as in the case of Model 2 was followed, with the only difference being the use of the OOB late fusion strategy in the weighted fusion step (instead of the proximity ratio scheme).
4. Model 4: The fourth model involved the same application of steps *A* and B from Model 2. Then, instead of retraining RF classifiers for the two modalities (as in Step *C* – Model 2) with the use of the final feature subsets, we opted to train Support Vector Machine (SVM) classification models. Finally, regarding the fusion step (step *D* – Model 2), we performed simple averaging of the probability scores provided by the SVM models. It should be noted that the probability scores of the SVM models were computed with the use of the Platt scaling method (Platt, 1999). Based on this technique, the output of a classification model is converted to a probability distribution over classes.
5. Model 5: In the case of the fifth model, steps A and B from Model 2 were applied in the same way (the only difference was the application of a different value/threshold for the Gini importance measure regarding the feature selection process in step B). Then, we applied early fusion to the resulting feature subsets from the two modalities (the feature subsets were concatenated into a common feature vector) and finally, a new RF model was trained with the use of the concatenated feature vector.

It should be noted that for Models 2-5, variables not specifically related to measurements from the left or right hemisphere (e.g. AGE, MMSE_bl, CSF, WM.hypointensities, etc.) were assigned to both modalities (left and right), before the feature selection process.

Finally, for the prediction of unknown cases based on the outputs of the ensemble’s models, a majority voting scheme was applied, meaning that the predicted class was the one that received the highest number of votes by the ensemble’s models. In the case of ties, the class with the highest probability estimate (provided by any of the models) was selected as the final prediction.

Our code for the experiments was written in R^3^. All RF models were developed using the randomForest^4^ package, while for the construction of the SVM models, the e1071^5^ package was used.

## 3 Experimental results

### 3.1 Experimental setup

With respect to the RF parameters that we used in the experiments, the number of trees for each RF model was empirically set (based on the OOB error estimate), while for each RF model and for each node split during the growing of a tree, the number *k* of the subset of variables used to determine the best split was set based on repeated 10-fold cross-validation that was performed using the caret^6^ package. Based on the aforementioned, the following parameter values were used for the RF models of the ensemble:

- Model 1: Number of trees = 2000, *k* = 53
- Models 2/3: Number of trees = 2000, 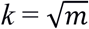 (where *m* is the total number of features)
- Model 5: Number of trees = 1000, *k* = 9

Regarding the threshold values for the Gini importance measure during the feature selection process in all RF models, the following values were used: **0.5** for Model 1, **0.75** for Models 2/3/4 and **4** for Model 5.

Finally, as already mentioned (Section 2.4.3), in Model 4, Support Vector Machine (SVM) classification models were trained for the two modalities/ final feature subsets. Specifically, for the left modality’s SVM model, a polynomial kernel was used, while a radial basis function (Gaussian) kernel was used for the right modality’s model. The aforementioned kernel types, as well as the parameter values for the two SVM models, were determined with the use of repeated 10-fold cross-validation (using the caret package). Specifically, for the left modality’s SVM model, the parameters *degree* and *scale* were set to 3 and 0.01, respectively, while for the right modality’s SVM model, the parameter *sigma* was set to 0.0163.

### 3.2 Selected Features Extraction

In Table 2, the selected features (with the use of the Gini importance measure) for each model of the ensemble are provided. We can notice that 53 features were selected for Model 1, 67 for Models 2/3/4, 41 for the left and right modality, respectively, and 9 features were selected in the context of Model 5.

**Table 2:** Selected features (based on the Gini importance measure) for each classification model of the ensemble

| Classification model | Selected features |
| --- | --- |
| **Model 1** | “AGE”, “MMSE_bl”, “Left.Inf.Lat.Vent”, “Left.Cerebellum.Cortex”, “Left.Hippocampus”, “CSF”, “Left.VentralDC”, “Left.vessel”, “Right.Inf.Lat.Vent”, “Right.Cerebellum.Cortex”, “Right.Hippocampus”, “Right.Amygdala”, “X5th.Ventricle”, “lh_lateralorbitofrontal_thickness”, “lh_medialorbitofrontal_thickness”, “lh_middletemporal_thickness”, “lh_parstriangularis_thickness”, “lh_posteriorcingulate_thickness”, “rh_entorhinal_thickness”, “rh_pericalcarine_thickness”, “rh_posteriorcingulate_thickness”, “rh_precentral_thickness”, “rh_insula_thickness”, “lh_medialorbitofrontal_area”, “lh_parsorbitalis_area”, “lh_temporalpole_area”, “rh_paracentral_area”, “rh_transversetemporal_area”, “lh_entorhinal_volume”, “lh_rostralanteriorcingulate_volume”, “rh_caudalanteriorcingulate_volume”, “rh_cuneus_volume”, “rh_entorhinal_volume”, “rh_rostralanteriorcingulate_volume”, “rh_transversetemporal_volume”, “lh_fusiform_thicknessstd”, “lh_parahippocampal_thicknessstd”, “lh_paracentral_thicknessstd”, “lh_posteriorcingulate_thicknessstd”, “rh_medialorbitofrontal_thicknessstd”, “rh_precentral_thicknessstd”, “rh_temporalpole_thicknessstd”, “rh_insula_thicknessstd”, “lh_frontalpole_meancurv”, “rh_lateraloccipital_meancurv”, “rh_medialorbitofrontal_meancurv”, “left_presubiculum”, “Right.Hippocampus_hipposubfields”, “left_CA1”, “right_presubiculum”, “right_CA2_3”, “right_subiculum”, “right_CA4_DG” |
| **Models 2/3/4** | **Left modality:** “AGE”, “MMSE_bl”, “Left.Lateral.Ventricle”, “Left.Inf.Lat.Vent”, “Left.Cerebellum.Cortex”, “Left.Pallidum”, “X3rd.Ventricle”, “Left.Hippocampus”, “Left.Amygdala”, “CSF”, “Left.Accumbens.area”, “Left.VentralDC”, “Left.vessel”, “Left.choroid.plexus”, “WM.hypointensities”, “non.WM.hypointensities”, “SubCortGrayVol”, “BrainSegVol.to.eTIV”, “MaskVol.to.eTIV”, “lh_bankssts_thickness”, “lh_caudalmiddlefrontal_thickness”, “lh_entorhinal_thickness”, “lh_inferiorparietal_thickness”, “lh_inferiortemporal_thickness”, “lh_isthmuscingulate_thickness”, “lh_lateraloccipital_thickness”, “lh_medialorbitofrontal_thickness”, “lh_middletemporal_thickness”, “lh_parahippocampal_thickness”, “lh_parstriangularis_thickness”, “lh_posteriorcingulate_thickness”, “lh_precuneus_thickness”, “lh_rostralanteriorcingulate_thickness”, “lh_superiorfrontal_thickness”, “lh_superiortemporal_thickness”, “lh_MeanThickness_thickness”, “lh_entorhinal_area”, “lh_inferiortemporal_area”, “lh_bankssts_volume”, “lh_entorhinal_volume”, “lh_fusiform_volume”, “lh_inferiorparietal_volume”, “lh_inferiortemporal_volume”, “lh_middletemporal_volume”, “lh_parahippocampal_volume”, “lh_supramarginal_volume”, “lh_bankssts_thicknessstd”, “lh_caudalmiddlefrontal_thicknessstd”, “lh_entorhinal_thicknessstd”, “lh_parahippocampal_thicknessstd”, “lh_paracentral_thicknessstd”, “lh_posteriorcingulate_thicknessstd”, “lh_rostralanteriorcingulate_thicknessstd”, “lh_superiorfrontal_thicknessstd”, “lh_insula_thicknessstd”, “lh_fusiform_meancurv”, “lh_inferiorparietal_meancurv”, “lh_inferiortemporal_meancurv”, “lh_medialorbitofrontal_meancurv”, “lh_frontalpole_meancurv”, “Left.Hippocampus_hipposubfields”, “left_presubiculum”, “left_CA1”, “left_CA2_3”, “left_fimbria”, “left_subiculum”, “left_CA4_DG” |
|  | **Right modality:** "AGE", "MMSE_bl", "CSF", "Right.Lateral.Ventricle", "Right.Inf.Lat.Vent", "Right.Cerebellum.White.Matter", "Right.Cerebellum.Cortex", "Right.Hippocampus", "Right.Amygdala", "rh_entorhinal_thickness", "rh_pericalcarine_thickness", "rh_posteriorcingulate_thickness", "rh_insula_thickness", "rh_paracentral_area", "rh_supramarginal_area", "rh_transversetemporal_area", "rh_caudalanteriorcingulate_volume", "rh_entorhinal_volume", "rh_rostralanteriorcingulate_volume", "rh_supramarginal_volume", "rh_transversetemporal_volume", "rh_isthmuscingulate_thicknessstd", "rh_medialorbitofrontal_thicknessstd", "rh_parahippocampal_thicknessstd", "rh_pericalcarine_thicknessstd", "rh_precentral_thicknessstd", "rh_temporalpole_thicknessstd", "rh_insula_thicknessstd", "rh_bankssts_meancurv", "rh_caudalanteriorcingulate_meancurv", "rh_inferiorparietal_meancurv", "rh_lateraloccipital_meancurv", "rh_medialorbitofrontal_meancurv", "rh_transversetemporal_meancurv", "Right.Hippocampus_hipposubfields", "right_presubiculum", "right_CA1", "right_CA2_3", "right_fimbria", "right_subiculum", "right_CA4_DG" |
| **Model 5** | "AGE", "MMSE_bl", "lh_medialorbitofrontal_thickness", "lh_parahippocampal_thicknessstd", "left_presubiculum", "rh_entorhinal_thickness", "rh_temporalpole_thicknessstd", "rh_lateraloccipital_meancurv", "right_subiculum" |

In Figure 3, boxplots for 5 features (for each diagnosis class) that were selected as important in all classification models of the ensemble are depicted:

**Figure 3:**
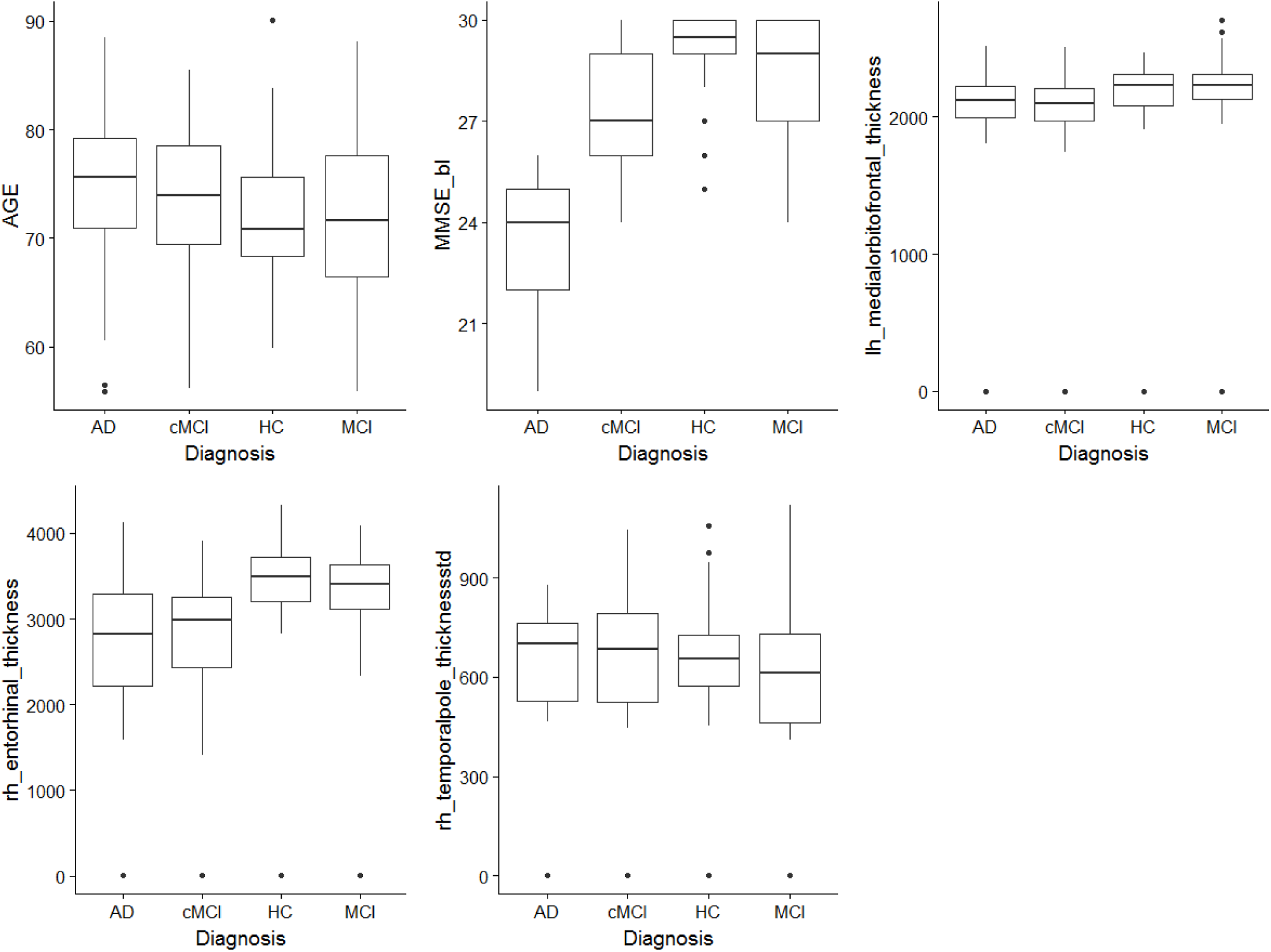
Boxplots for 5 features (for each diagnosis class), selected as important in all models of the ensemble

The confusion matrix for the predictions of the 160 test set subjects (without the 340 dummy subjects) can be seen in Table 3, while in Table 4, more detailed results (in terms of precision, recall and F-score measures for each class, their corresponding macro-averaged values and accuracy) with respect to the ensemble’s performance on the test set are provided:

**Table 3:** Confusion matrix for the predictions of the 160 subjects of the test set (without the 340 dummy test subjects)

| Class | HC predicted | MCI predicted | AD predicted | cMCI predicted |
| --- | --- | --- | --- | --- |
| HC real | 24 | 9 | 1 | 6 |
| MCI real | 14 | 14 | 7 | 5 |
| AD real | 0 | 0 | 38 | 2 |
| cMCI real | 5 | 8 | 4 | 23 |

**Table 4:** Test set results

| Class | Precision | Recall | F-score |
| --- | --- | --- | --- |
| HC | 55.8% | 60.0% | 57.8% |
| MCI | 45.1% | 35.0% | 39.4% |
| AD | 76.0% | 95.0% | 84.4% |
| cMCI | 63.8% | 57.5% | 60.5% |
| **Macro-average** | **60.2%** | **61.9%** | **60.5%** |
| **Accuracy** | **61.9%** |  |  |

From the results in Table 4, we notice that our classification ensemble achieves an **accuracy of 61.9% (the best performance achieved in the neuroimaging challenge)**, as well as macro-averaged values of 60.2%, 61.9% and 60.5% for the precision, recall and F-score measures, respectively. The best performance is achieved for the class “AD” (precision 76.0%, recall 95.0%, F-score 84.4%), while the worst results are attained for the “MCI” class (precision 45.1%, recall 35.0% and F-score 39.4%).

## Discussion

In the present study, we managed to achieve a high level of classification accuracy in a blind dataset working for the first time in a four-class AD-based problem. In the feature space, we added morphological MRI-based features that in recent years have been shown to increase classification accuracy for the automatic diagnosis of AD, such as cortical thickness, subcortical volumes and hippocampal subfields (Desikan et al., 2009; Vasta et al., 2016; de Vos et al., 2016). Regarding the classification strategy, we adopted an RF approach, designing various models for a better learning of the feature space in the internal dataset, thus improving the generalization of the whole model. Then, we performed classification using the selected feature set from the training set to the blind testing dataset. We achieved a 61.9% classification performance for the simultaneously discrimination of four groups (HC, MCI, cMCI and AD). It is the very first time in the literature where classification is performed simultaneously in a four AD-based class problem using MRI.

Regarding the modalities, many studies focusing on predictive biomarkers for either AD or MCI or both conditions investigated structural MRI data alone (Lebedev et al., 2014; Moradi et al., 2015; Nanni et al., 2016; Salvatore et al., 2015,2016; Ardekani et al., 2017; Lebedeva et al., 2017) or combined with features extracted from other modalities, like FDG-PET (Gray et al., 2013; Sivapriya et al., 2015), florbetapir-PET (Wang et al., 2016), FLAIR (Oppedal et al., 2015) and fMRI (Tripoliti et al., 2007; Son et al., 2017).

Focusing on the cohort diagnosis and the targeted groups, two studies (Tripoliti et al., 2007; Lebedev et al., 2014) investigated Alzheimer’s patients (AD) and healthy controls (HC), four studies (Cabral et al., 2013; Sivapriya et al., 2015; Maggipinto et al., 2017; Son et al., 2017) examined AD, HC and Mild Cognitive Impairment (MCI), two studies (Gray et al., 2013; Moradi et al., 2015) considered AD, HC, stable MCI (sMCI) and progressive MCI (pMCI, converted to AD), two had sMCI and pMCI (Wang et al., 2016; Ardekani et al., 2017), one had HC and MCI (Lebedeva et al., 2017) and one (Oppedal et al., 2015) had AD, HC and Lewy-body dementia (LBD) patients.

A total of eight studies (Tripoliti et al., 2007; Cabral et al., 2013; Lebedev et al., 2014; Moradi et al., 2015; Sivapriya et al., 2015; Ardekani et al., 2017; Lebedeva et al., 2017; Maggipinto et al., 2017) applied a feature selection strategy for reducing the dimension of the variables space. In two out of eight cases, the number of trees used in the RF was not specified (Moradi et al., 2015, Son et al., 2017).

Finally, the reported classification accuracies for binary classifiers were: 64.63% for HC-AD (Cabral et al., 2013), HC-AD: 89% / HC-MCI: 74.6% / sMCI-pMCI: 58.4% (Gray et al., 2013), HC-AD: 90.3% (Lebedev et al., 2014) and sMCI-pMCI: 82% (Moradi et al., 2015). Two studies that adopted multi-class classifiers reported 96.3% classification accuracy for HC-MCI-AD: (Sivapriya et al., 2015) and 87% for HC-LBD-AD (Oppedal et al., 2015). However, both of them used multimodal features, in contrast to only MRI-based studies.

A recent multi-class study based on MRI reported a classification accuracy of ∼60% for HC-MCI-AD using a regularized extreme learning machine and PCA for feature selection (Lama et al., 2017). The whole approach was based on an internal cross-validation scheme without attempting to classify a second external blind dataset. In our study, we outperformed the best reported single-modality classification performance for HC-MCI-AD focusing on the distinction of MCI to cMIC-MCI.

In this study, we applied early fusion as well as late fusion strategies based on RF’s operational features, namely the OOB error and proximity ratios. For the prediction of an unknown case, the RF models provide probability estimates per class for that case based on a weighting fusion strategy (Liparas et al., 2014). We built in total five models by splitting the feature space in the left and right hemispheres. Finally, for the prediction of unknown cases based on the outputs of the ensemble’s models, a majority voting scheme was applied, meaning that the predicted class was the one that received the highest number of votes by the ensemble’s models. Finally, the class with the highest probability estimate (provided by any of the models) was selected as the final prediction. This is the very first time that such an RF-based scheme was performed and particularly an automatic multi-class classification scheme tailored to Alzheimer’s disease and structural MRI modality.

The most discriminative structural features were the following: age, MMSE scores, the thickness of the right entorhinal thickness, right temporal pole thickness and the right medial orbitofrontal cortex thickness. Right entorhinal atrophy has been revealed as a consequence of frontotemporal dementia and Alzheimer’s disease (Frisoni et al., 1999). Thickness of the right temporal pole has been linked to the lateralization effect of semantic dementia (Kumfor et al., 2016) while the thickness of the right medial orbitofrontal cortex is a key brain area that differentiates the prodromal stage of AD from normal aging (Blanc et al., 2015).

It is important to underline here that both training and testing datasets were age-matched. We employed age as a possible feature among the MMSE and MRI-based on the assumption that the synergy with morphological properties could play a key role in the improvement of classification accuracy. A recent study explored the synergy of age and APOE for predicting progression from MCI to AD (Korolev et al., 2017). Another study demonstrated a Bayesian model for the early prediction and early diagnosis of AD (Alexiou et al., 2017). We hypothesize that age will have a weaker relation to a prediction model, e.g. for the conversion of MCI to AD for a group, following a physical and cognitive intervention.

RF has been successfully applied to a wide range of disciplines and several studies that make use of RF in the neuroscience domain can be mentioned. For instance, Ramirez et al. (2010) presented a computer aided diagnosis (CAD) method for the early detection of the Alzheimer’s disease (AD), based on partial least square (PLS) regression for feature extraction and RF for single photon emission computed tomography (SPECT) image classification. The experimental results of their study showed that the proposed PLS-RF system’s generalization error converges to a limit as the number of trees in the RF model increases and is affected by the strength of the trees in the model, as well as the correlation between them. In another study, Smith et al. (2013) performed prediction of the concentrations of 9 neurochemicals in the vestibular nucleus complex and cerebellum by means of Random Forest regression (RFR) and compared the results with those of multiple linear regression (MLR). In general, the experimental results demonstrated the superiority of MLR over RFR in terms of predictive value and error. Nevertheless, an interesting conclusion of the study was that RFR can still have good predictive value in certain cases. Lebedev et al. (2014) investigated the effectiveness of RF classifier ensembles in the detection and prediction of AD in terms of accuracy and between-cohort robustness. The ensembles were trained with the use of different structural MRI measures and they resulted in significantly better classification performance compared to the reference model (linear Support Vector Machine). Finally, McKinley et al. (2016) proposed a method, called fully automated stroke tissue estimation using random forest classifiers (FASTER), which estimates the penumbra (tissue-at-risk) volume in the context of ischemic stroke treatment. The method utilizes multimodal MRI in order to predict tissue damage in the case of persistent occlusion, as well as of complete recanalization.

A recent systematic review of RF algorithms tailored to the classification of neuroimaging data in AD underlines the limitations of single modalities, the best accuracies of multimodal imaging and overfitting (Sarica et al., 2017). Finally, they suggested the need for the use of machine learning techniques for the early prediction of the progression from MCI to AD.

Complementary to the aforementioned structural features, hippocampal volume has been listed high in the ranking of features. Hippocampal volume plays a key role in early dementia and cognitive decline. Hippocampal atrophy is higher in AD compared to MCI and healthy controls (Heiyer el al., 2010). Hippocampal volumes were also inversely correlated with age in older healthy controls while in Alzheimer’s disease hippocampal atrophy in the body and tail of overlap with atrophy was also observed in healthy controls. In contrast, the atrophy in the anterior and dorsal CA1 subfield involved in Alzheimer’s disease was not found in normal ageing (Frisoni et al., 2008). Parcellating the hippocampus with Freesurfer 6.0 will increase the distinction of atrophy between healthy control and Alzheimer’s disease patients and also in mild cognitive impairment subgroups (Iglesias et al., 2015).

## Limitations of the Study

In the current study, we attempted to predict the labels of an unknown dataset in a four-class problem. We achieved a classification accuracy of 61.9%, which is low for a classification performance, especially for AD, but the best till now in the literature. This open competition with a common starting point for every team underlined the limitations of a single imaging modality in the construction of a reliable biomarker that can track every pre-stage of AD and distinguishes MCI from cMCI. It is vital in the near future to combine features from multimodal imaging with genetic risk for AD (Foley et al.,2017), various neuropsychological estimates and also complementary features, such as living habits (Alexiou et al., 2017), for the design of a better early diagnostic model for Alzheimer’s disease. To reveal the complementary information shared between every group of features in a final model and also their causal role, accelerated longitudinal studies are very important (Teipel et al., 2015). We strongly believe that the current methodology could be a substrate to fuse multimodal features and to further predict the clinical status of an unknown dataset.

In the future, we will attempt to use the same methodological approach, focusing also on subjects with a longer follow-up period with main scope to improve the sensitivity of our algorithm in discriminating stable vs progressive MCI subjects (Lebedev et al., 2014). In addition, we will extract features from static and dynamic functional brain networks, based on resting-state fMRI recordings for building multi-modal biomarkers.

## Conclusions

Our methodology based on RF and structural MRI features produces the highest classification accuracy for a multi-class AD-based problem. It is the very first study that attempted to simultaneously classify four classes (HC, cMCI, MCI, AD), and achieved a classification accuracy of 61.9% in a blind external validation dataset. Our approach could be useful also for multimodal biomarkers focusing on novel and robust AD biomarkers.

## Footnotes

2 The preprocessing of the T1-weighted Magnetic Resonance Images (MRI) was realized by the organizers of the competition. Information can be found here: https://inclass.kaggle.com/c/mci-prediction

2 http://adni.loni.usc.edu/methods/mri-analysis/mri-acquisition/

3 https://www.r-project.org/

4 https://cran.r-project.org/web/packages/randomForest/index.html

5 https://cran.r-project.org/web/packages/e1071/index.html

6 https://cran.r-project.org/web/packages/caret/index.html

## Acknowledgement

We would like to thank the anonymous reviewers for their valuable comments that further improved the quality of the manuscript. SID was supported by a MRC grant MR/K004360/1 (Behavioural and Neurophysiological Effects of Schizophrenia Risk Genes: A Multi-locus, Pathway Based Approach). SID is also supported by a MARIE-CURIE COFUND EU-UK Research Fellowship.

